# Rigorous (in)validation of ecological models

**DOI:** 10.1101/2024.09.19.613075

**Authors:** Chuliang Song, Jonathan M. Levine

## Abstract

The complexity of ecosystems poses a formidable challenge for confidently invalidating ecological models, as current practices struggle to distinguish model inadequacies from the confounding effects of unobserved biotic or abiotic factors. The prevailing inability to falsify models has resulted in an accumulation of models but not an accumulation of confidence. Here, we introduce a new approach rooted in queueing theory, termed the *covariance criteria*, that establishes a rigorous test for model validity based on covariance relationships between observable quantities. These criteria set a high bar for models to pass by specifying necessary conditions that must hold regardless of unobserved factors. We demonstrate the broad applicability and discriminatory power of the covariance criteria by applying them to three long-standing challenges in ecological theory: resolving competing models of predator-prey functional responses, disentangling ecological and evolutionary dynamics in systems with rapid evolution, and detecting the often-elusive influence of higher-order species interactions. Across these diverse case studies, the covariance criteria consistently rule out inadequate models, while building strong confidence in those that provide strategically useful approximations. The covariance criteria approach is mathematically rigorous, computationally efficient, and often non-parametric, making it immediately applicable to existing data and models.

---

Ecology textbooks grow ever longer, because older theories stick around even as new ones pile up.

— Stefano Allesina

## 1 Complexity in invalidating ecological model

Population abundance is the ever-present variable in the equation of life on Earth. To decipher the drivers behind the fluctuations of population abundance, ecologists construct mathematical models—simplified representations that capture an ecosystem’s key dynamics while making judicious sacrifices of nature’s full complexity. This synergy between data and modeling forms the foundation of contemporary ecology (Kingsland, 1991; Grainger et al., 2022). Yet, this endeavor faces a fundamental challenge: how can we confidently adjudicate which models provide useful approximations of nature, and which are oversimplified caricatures? The stark reality is that even for predator-prey interactions, there exist more than 40 distinct models of how predator feeding rate depend on prey abundance (reviewed in Novak and Stouffer 2021). This plethora of alternatives stems from the prevailing inability, using conventional practices, to decisively validate some models and invalidate others against empirical data.

To illustrate the limitations of current approaches, consider a textbook example of the coupled population dynamics of snowshoe hares and Canadian lynx in boreal forests (Leigh, 1968) (Figure 1A). The dynamics is classically modelled using the Lotka-Volterra (LV) predator-prey model (Figure 1B). A common validation approach is to compare the qualitative behaviors between data and model prediction (Figure 1C). In this example, the Lotka-Volterra model predicts coupled cycles of predator and prey abundances with a fixed amplitude and period length—the hallmark of the “predation cycle” in graphical predator-prey theory (Volterra, 1926; Rosenzweig and MacArthur, 1963). The qualitative resemblance of the data to these predicted cycles provides some confidence in the model’s validity. Another common approach is fitting models to data and assessing goodness-of-fit or forecasting power (Figure 1D). For this example, the predator-prey dynamics can be approximated with a given set of parameters in LV dynamics, providing support for the proposed model. These two approaches represent the mainstream for validating models against time series data.

**Figure 1.**
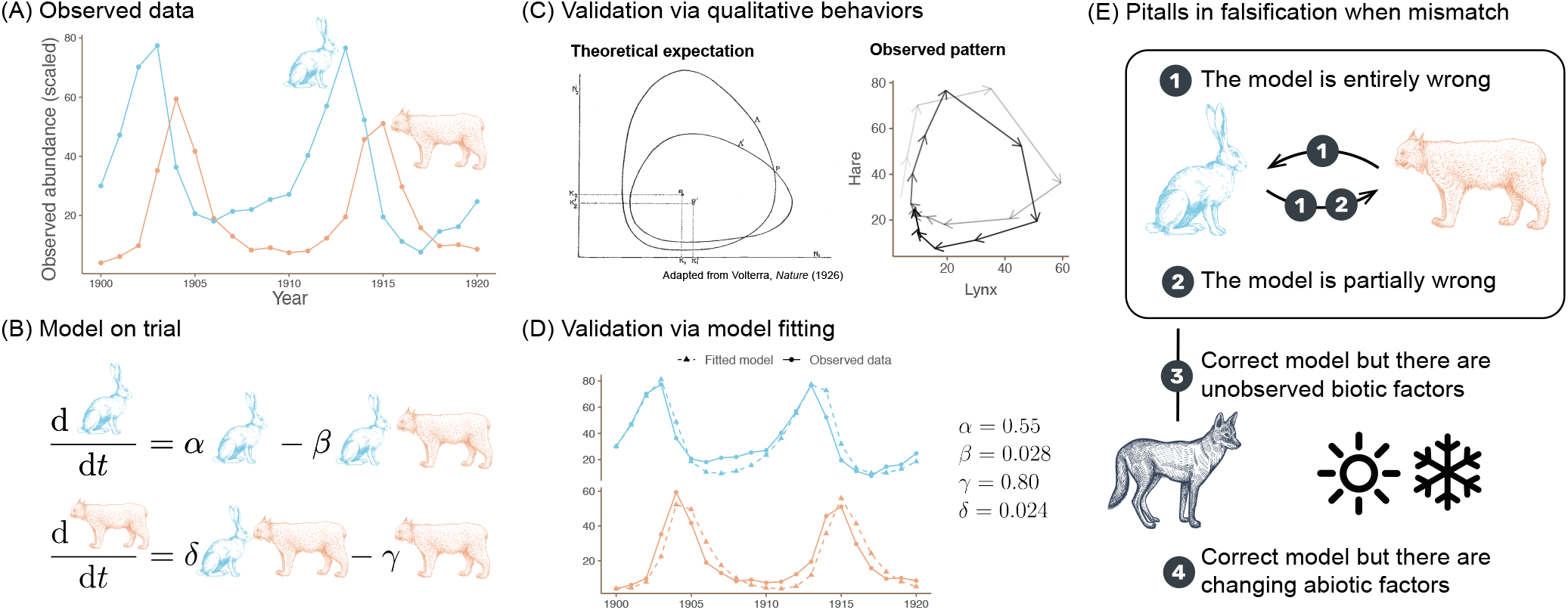
Common approaches in ecological model validation and limitations. Panels (A) and (B) depict a classic example of predator-prey dynamics: the population fluctuations of snowshoe hares (blue) and Canada lynx (red), and the Lotka-Volterra (LV) model as a candidate to describe the underlying processes. Panels (C) and (D) demonstrate two validation approaches: comparing qualitative behaviors (e.g., cycles in both data and model), and fitting the model to the data to examine its explanatory or predictive power. Panel (E) highlights challenges in interpreting validation results. A mismatch between a model and data does not necessarily prove the model is entirely incorrect (case 1), as the discrepancy could stem from the model being partially incorrect (case 2), unobserved biotic interactions (case 3), or abiotic influences (case 4). Current methods often cannot decisively determine which of these cases is responsible for the mismatch.

But what if the model and data diverge? Does that mean the model is *invalidated*? Returning to the hare-lynx example, over longer time scales than shown in Figure 1, the observed population cycles diverge from the classic Lotka-Volterra predictions: in fact, the data suggest a “reversed cycle,” implying the nonsensible result that hares eat lynx (Gilpin, 1973). However, interpreting such discrepancies is fraught with ambiguity. One possibility is that these deviations in qualitative patterns invalidate the core assumptions of the Lotka-Volterra framework, demanding an alternative theoretical model for the entire predator-prey system (Cortez and Weitz 2014; Case ❶ in Figure 1E). Alternatively, the model could accurately describe the dynamics of one species (e.g. lynx) while failing for the other (e.g. hares) due to missing factors specific to that species (Stenseth et al. 1997; Case ❷ in Figure 1E). In contrast, it is also possible that the model correctly describes the lynx-hare interaction, but fails to include all of the other variables driving the observed dynamics. These could be other uncontrolled biotic factors (Case ❸ in Figure 1E) such as the hare-vegetation interaction (Blasius, Huppert, and Stone, 1999) or disease epidemics. Or it could be that there are other uncontrolled abiotic factors (Case ❹ in Figure 1E), such as environmental fluctuations altering species parameters over time (Hone, Krebs, and O’Donoghue, 2011; Yan et al., 2013; King and Schaffer, 2001). In sum, current practices make it challenging to judge whether the model is truly valid, partially valid, or simply wrong.

This long-standing challenge of model validation has plagued ecology, leaving the true scope of even classic models like the Lotka-Volterra formulation unresolved. Here, we address this fundamental problem in ecological modeling by introducing a method originally developed by biophysicists (Hilfinger, Norman, Vinnicombe, et al., 2016). In essence, this method uncovers the (mostly unique) inherent structure of temporal covariance between model elements, a constraint that remains invariant regardless of unknown ecological factors. By leveraging this inherent constraint, we can make strong statements about model validity. In the following sections, we first introduce the theoretical foundations of this approach. We then demonstrate its discriminatory power by applying it to three key problems in ecology: resolving debates on the functional forms of predator-prey interactions, disentangling the interplay of ecology and rapid evolution, and detecting signals of higher-order species interactions. Through these case studies, we illustrate how this rigorous test of model validity can decisively invalidate flawed models, build confidence in those that provide useful approximations, and guide the development of more robust ecological theory.

### 2 Covariance criteria for model (in)validation

In this section, we introduce the theoretical framework for the covariance criteria approach and demonstrate its application to ecological models and data. We start by presenting the core concepts and mathematical foundations. We then illustrate how to apply the framework to a simple work-out example with statistical methods. Finally, we discuss the advantages of this approach over current model validation practices.

### 2.1 General theoretical framework

The fluctuations in population abundances that we observe in nature arise from a fundamental imbalance between two opposing forces: the *gain rate*, which encompasses processes that increase population size (e.g., births, immigration, mutualism), and the *loss rate*, which includes processes that decrease it (e.g., deaths, emigration, competition). In general, ecological models describing the dynamics of population abundance can be partitioned in the following form:

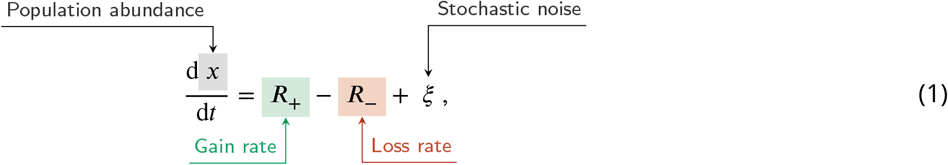

where *R*_+_ is the gain rate, *R*___ is the loss rate, and ξ is the stochastic noise. The gain rate *R*_+_ and loss rate *R*___ can be complex functions of both biotic (e.g., interactions with other species) and abiotic (e.g., environmental) factors. Any meaningful ecological model must include this partitioning of gain and loss; otherwise, the model would predict either indefinite growth or inevitable extinction. We can illustrate this partitioning using the predator dynamics from the Lotka-Volterra (LV) model:

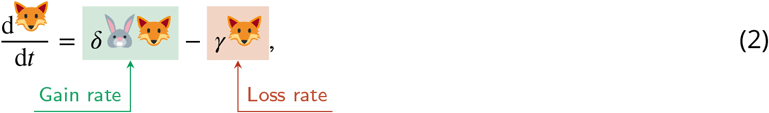

where 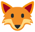 denotes the predator and 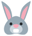 denotes the prey.

When gain and loss rates are perfectly counterbalanced, the population maintains a steady state, or equilibrium. However, such perfect balance is seldom encountered in the real world. Instead, we witness periods where gains outweighs losses, leading to population growth, interspersed with periods where losses dominate, causing population decline. This constant interplay between gains and losses generates the dynamic fluctuations in abundance that characterize most natural populations.

While it is intuitive that changes in population abundance relate to the gain-loss imbalance, mathematically quantifying this relationship is challenging. The challenge arises from the fact that the gain and loss imbalance is related to the *rate of change* in abundance, not the *raw* abundance. To resolve this, we can use the concept of covariance, a statistical measure of how two variables change together. Consider a scenario where gain rates tend to be higher when the population abundance is high. We can say that the gain rate and abundance covary *positively*. Importantly, in a system that fluctuates but avoids unchecked growth, the loss rate must also increase when abundance is high to counteract the increased gain. Thus, the loss rate and population abundance must also covary *positively*.

Formally, this constraint, while grounded in sophisticated mathematics, is captured in a surprisingly simple equality (Hilfinger, Norman, Vinnicombe, et al., 2016):

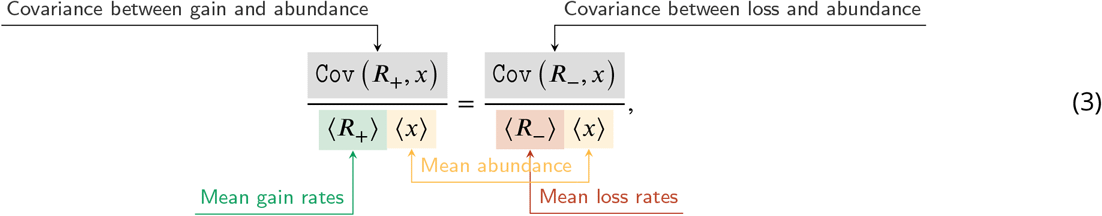

where `Cov` denotes covariance and ⟨ ⟩ denotes mean (Figure 2A). In words, the equality (Eq. 3) essentially states that the normalized covariance between gain rate and abundance is mirrored by the normalized covariance between loss rate and abundance.

**Figure 2.**
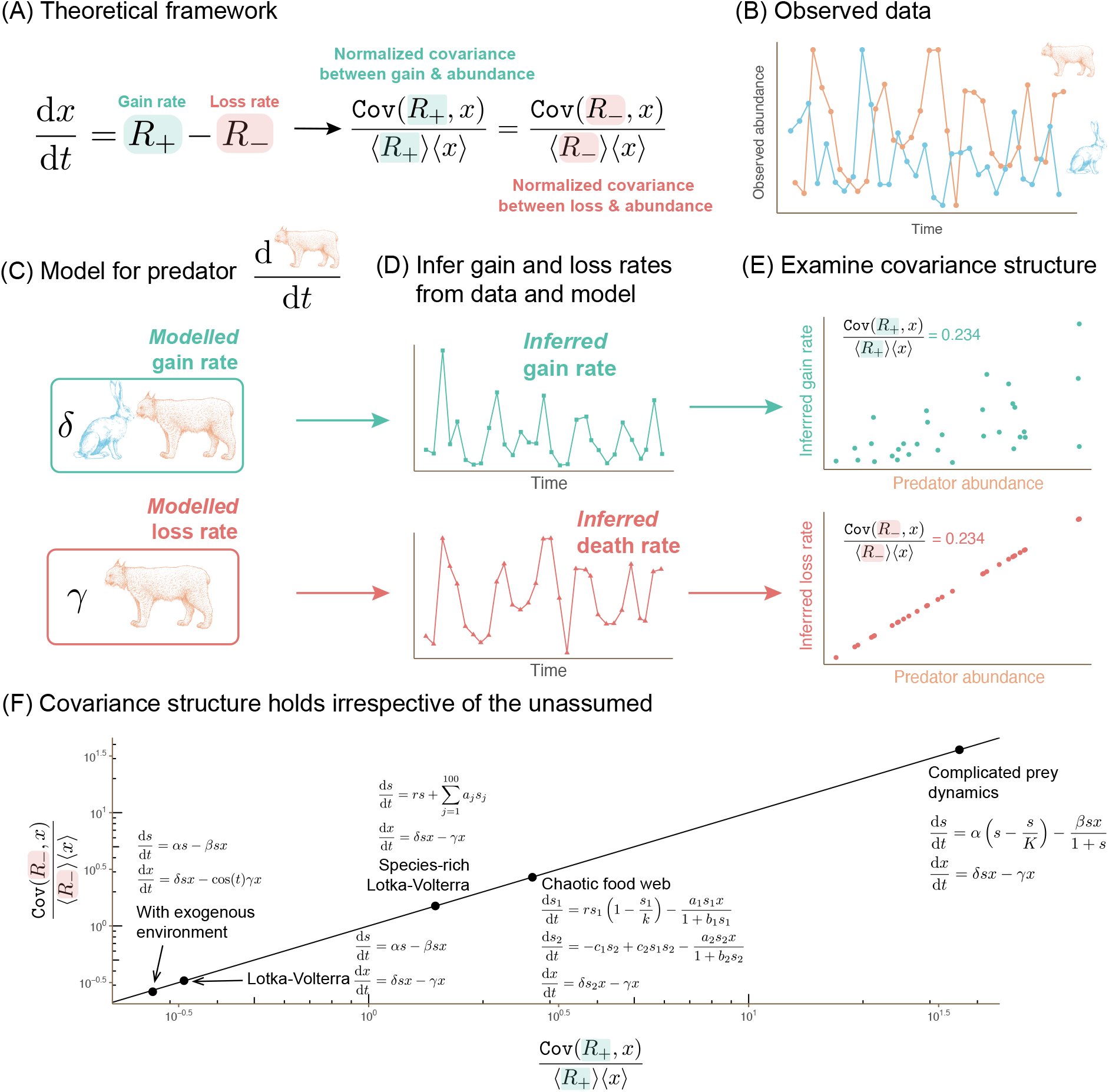
Covariance criteria for model validation. Panel (A) presents the general principle of covariance criteria: if a model accurately captures the underlying processes, the normalized covariance between the gain rate and abundance should equal the normalized covariance between the loss rate and abundance. Panels (B) to (E) illustrate the workflow of this approach to assess whether the Lotka-Volterra (LV) model can describe predator dynamics in a specific system. Panel (B) shows a subset of the predator-prey data (Blasius, Rudolf, et al., 2020). Panel (C) shows demonstrates how the LV model partitions predator dynamics into gain rate (terms causing abundance increase) and loss rate (terms causing abundance decrease). Panel (D) integrates the modeled gain and loss rates with the observed data to infer the empirical gain and loss rates across time. Panel (E) calculates the normalized covariances between the inferred gain/loss rates and predator abundance. If these covariances are equivalent, it strengthens confidence in the model’s validity. Conversely, significant discrepancies indicate the LV model’s inadequacy in describing the predator’s dynamics. Panel (F) emphasizes the universality of the covariance criteria (see details in Appendix A). This means that the criteria hold true even if the model does not explicitly include all factors influencing the system. To illustrate this, simulations are used where the predator follows the LV model, but the rest of the ecological community can exhibit arbitrarily complex dynamics. These simulations serve purely as an illustrative aid, as the method’s core strength lies in its mathematical rigor.

Mathematically, this equality is known as the second order moment equation derived from Little’s law in queuing theory (Little, 1961; Little, 2011). Note that our representation is slightly different from Hilfinger, Norman, Vinnicombe, et al. (2016) as we use a continuous stochastic dynamic instead of discrete one. We do so because continuous models are more commonly used in ecology, and also because population abundances are measured as density or biomass that do not have a direct discrete interpretation.

### 2.2 An illustrated worked-out example

This covariance structure serves as a simple test to validate or invalidate a model. If the model accurately reflects the observed ecological dynamics, the equality (Equation 3) will hold, and the model passes the test. If the data and constraint don’t match, the equality won’t hold, and the model is falsified. We call this approach the *covariance criteria*.

We next illustrate how the covariance criteria works in practice using the predator dynamics from the Lotka-Volterra (LV) model (Eqn. 2). The model defines the gain rate as proportional to the product of prey and predator abundances (representing successful predation leading to reproduction), while the loss rate is proportional to predator abundance alone (representing mortality). Applying the general covariance constraint (Equation 3) to this model, we get:

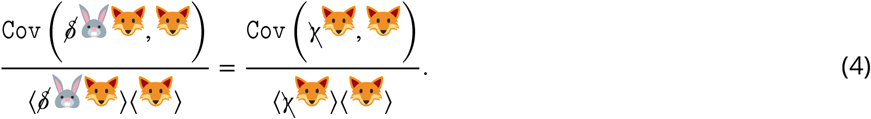

Since the mean and covariance operators are linear, we can simplify this further by canceling out the constant parameters δ and γ:

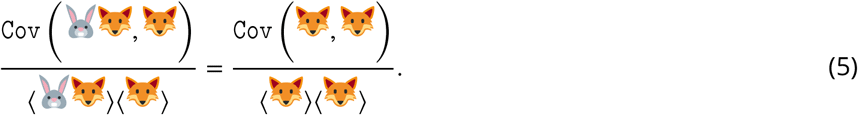

This simplification is important. It lets us evaluate the covariance criterion directly from the observed abundance data *without* needing to estimate the often unknown parameter values, making it a non-parametric test.

We next test this model using a subset of data from a planktonic predator-prey system (Blasius, Rudolf, et al., 2020) (Figure 2B). The data include time series measurements of both predator and prey abundance. Although we can’t directly observe the gain and loss rates in the data, the model allows us to infer them at each time point based on the observed abundance data (Figure 2C). For instance, the gain rate is inferred as the product of prey and predator abundance at each time point (recall that the parameter in front of this product cancels out in Equation 5).

The covariance criteria then examine how these inferred gain and loss rates covary with the observed predator abundance (Figure 2C). The scatter plot of inferred gain rate against predator abundance shows more scatter compared to the plot of inferred loss rate against predator abundance. This is expected because the predator’s gain rate depends on both prey and predator abundance, while the loss rate depends solely on predator abundance. However, it is important to remember that the covariance criteria rely on the calculated covariance values to provide the quantitative measure of the relationship between the rates and predator abundance, not the visual spread of the scatter plots. In this specific example, both normalized covariances turn out to be 0.234. This suggests that the LV model, in this case, aligns with the covariance criterion and adequately explains the observed predator dynamics.

### 2.3 Addressing noise and limited data

Ecological time series data is often noisy and limited in length, which can make it difficult to draw reliable inferences from point estimates of covariance values alone. To estimate uncertainty around the covariance measures, we can use bootstrapping. We repeatedly draw random samples from the original time series data with replacement and recalculate the gain and loss rate covariances for each resampled dataset. This generates distributions of covariance that capture the inherent variability in the data.

We then compare the distribution of gain rate covariances and the distribution of loss rate covariances. To assess the statistical significance of any observed difference between these distributions, we examine the distribution of their pairwise differences. A significant overlap between the pairwise difference distribution and zero suggests that the model-predicted equality between the covariances is statistically supported. In contrast, a pairwise difference distribution that is clearly shifted away from zero provides strong evidence that the model violates the covariance criteria. To quantify this difference, we calculate Cohen’s d, a standard measure of effect size between the pairwise difference distribution and zero. A z-score below a threshold (typically 1.96 for 95% confidence) indicates the distributions are statistically indistinguishable—the covariances are not different from each other, and the model passes the test. This threshold can be adjusted to control the balance between false positives and false negatives as needed. For the planktonic predator prey example in Figure 2, the z-score is 0.04, indicating that the two covariances are likely the same.

We have developed the R package ecoModelOracle to streamline this statistical analysis, making it easier for users to implement the approach.

### 2.4 Applicability of the covariance criteria

The covariance criteria works under remarkably broad conditions, thanks to the generality of Little’s Law in queuing theory (Little, 1961; Little, 2011). It applies rigorously to stationary systems, where long-term statistical patterns remain constant over time, regardless of whether they follow typical Markovian dynamics (where the future depends only on the present, as most ecological models do) or more complex non-Markovian dynamics (where the ecosystem’s history influences its future, as in the presence of time delays). Moreover, the criteria also holds for some non-stationary dynamics, like cyclo-stationary systems (where statistical patterns repeat predictably, as with seasons). One might assume that such broad applicability renders the criteria a mere abstract principle with limited practical utility. Surprisingly, as we show here, it imposes a stringent test for models to pass. When the biophysicists who pioneered this method applied it to gene expression data, nearly all published models failed to meet the criteria (Hilfinger, Norman, and Paulsson, 2016). This means that when a model *does* pass the test, we can have strong confidence in its validity.

Due to its generality, the covariance criteria can be used to interrogate models in a more precise manner than traditional approaches (Figure 1E). First, the criteria directly test the dynamics of individual species, eliminating the need to know whether the model is correct for the entire system (Case ❶ vs. Case ❷ in Figure 1E). For instance, we can validate whether the predator dynamics follow the LV model regardless of whether the prey dynamics also follow the LV model (‘Lotka-Volterra’ in Figure 2F) or follow a more complex model (‘complicated prey dynamics’ in Figure 2F). Second, the covariance test is invariant to unknown species that indirectly interact with the species under examination (Case ❸ in Figure 1E). For instance, the test for the predator remains valid even when we add many other species that only interact with the prey (‘Species-rich Lotka-Volterra’ in Figure 2F), including interactions that ultimately drive chaotic dynamics (‘Chaotic food web’ in Figure 2F). Lastly, the covariance criteria can sometimes tolerate unknown abiotic factors (case ❹ in Figure 1E), particularly when the environment acts as an exogenous driver statistically independent of population abundance (‘With exogenous environment’ in Figure 2F).

To illustrate the power of the approach, we revisit the long-standing debate on whether the hare-lynx dynamics in the Canadian boreal zone adhere to the Lotka-Volterra model. We do so by applying the covariance criteria to the full dataset from the system. For hares, the calculated z-score between the distributions of gain and loss covariances is 4.9, a value that strongly suggests unequal covariances and the Lotka-Volterra model’s inadequacy in capturing hare dynamics. In contrast, the z-score for the gain and loss covariance distributions for lynx is 1.7. This implies that one cannot statistically reject the equality of covariances, supporting the Lotka-Volterra model as a potentially useful approximation for lynx dynamics. These findings mirror Case ❷ in Figure 1, aligning with the hypothesis proposed by Stenseth et al. (Stenseth et al., 1997) that the LV model might be valid for lynx but not for hares in this predator-prey system.

## 3 Three Case Studies

In this section, we apply the covariance criteria to tackle three long-standing problems in ecology. We begin by investigating the fundamental nature of predation, using the criteria to rigorously test different functional forms describing the predator-prey interaction. We then study the integration of rapid evolutionary processes into ecological models, leveraging the criteria to potentially pinpoint where evolutionary forces significantly shape species dynamics. Finally, we search for the often-hidden influence of higher-order interactions within ecosystems, harnessing the criteria to uncover complex relationships that extend beyond simple pairwise effects.

### 3.1 Reverse engineering the nature of predation

Predation is a key force structuring ecological communities (Pringle et al., 2019). Yet, modeling this interaction remains a challenge. A central debate persists over whether predation is best described by Lotka-Volterra (in which predation is a function of the product of predator and prey abundances), or instead, additionally dependent on the ratio of predator and prey individuals in the system. This debate has persisted for decades (Abrams and Ginzburg, 2000; Arditi and Ginzburg, 2012; Abrams, 2015; Tyutyunov and Titova, 2020; Ginzburg and Damuth, 2022), partly because traditional methods often struggle to definitively rule out alternative explanations (Morin, 2009). Here, we demonstrate how the covariance criteria can help resolve this long-standing question.

We examine three possible prey dynamics: LV dynamics with and without self-regulation, and ratio-dependent dynamics (Akcakaya, Arditi, and Ginzburg, 1995; Arditi and Ginzburg, 1989),

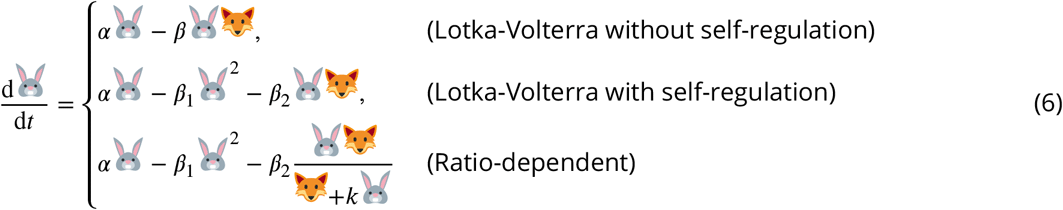

Similarly, we consider three possible predator dynamics:

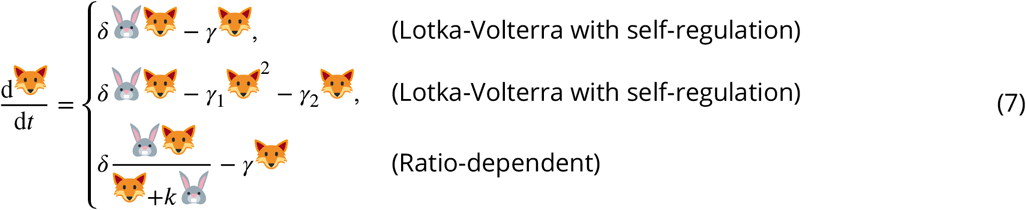

To rigorously test these models, we analyze a unique long-term dataset with replicated predator-prey dynamics under various conditions (Blasius, Rudolf, et al., 2020). The dataset tracks the relationship between the aquatic invertebrate consumer *Brachionus calyciflorus* and its green algae prey *Monoraphidium minutum*. We find that the prey dynamics align most closely with LV dynamics with self-regulation (Figure 3A-C), while the predator dynamics align most closely LV dynamics without self-regulation (Figure 3D-F). See Figure S2 for the z-score statistical test for each model. For both species, the model with ratio dependent interactions deviates most from the equality constraint posed by the covariance criteria. Our findings therefore provide compelling evidence that predation in this system is prey-dependent as posed in the Lotka-Volterra model, and not a function of the ratio of predators and prey in this system.

**Figure 3.**
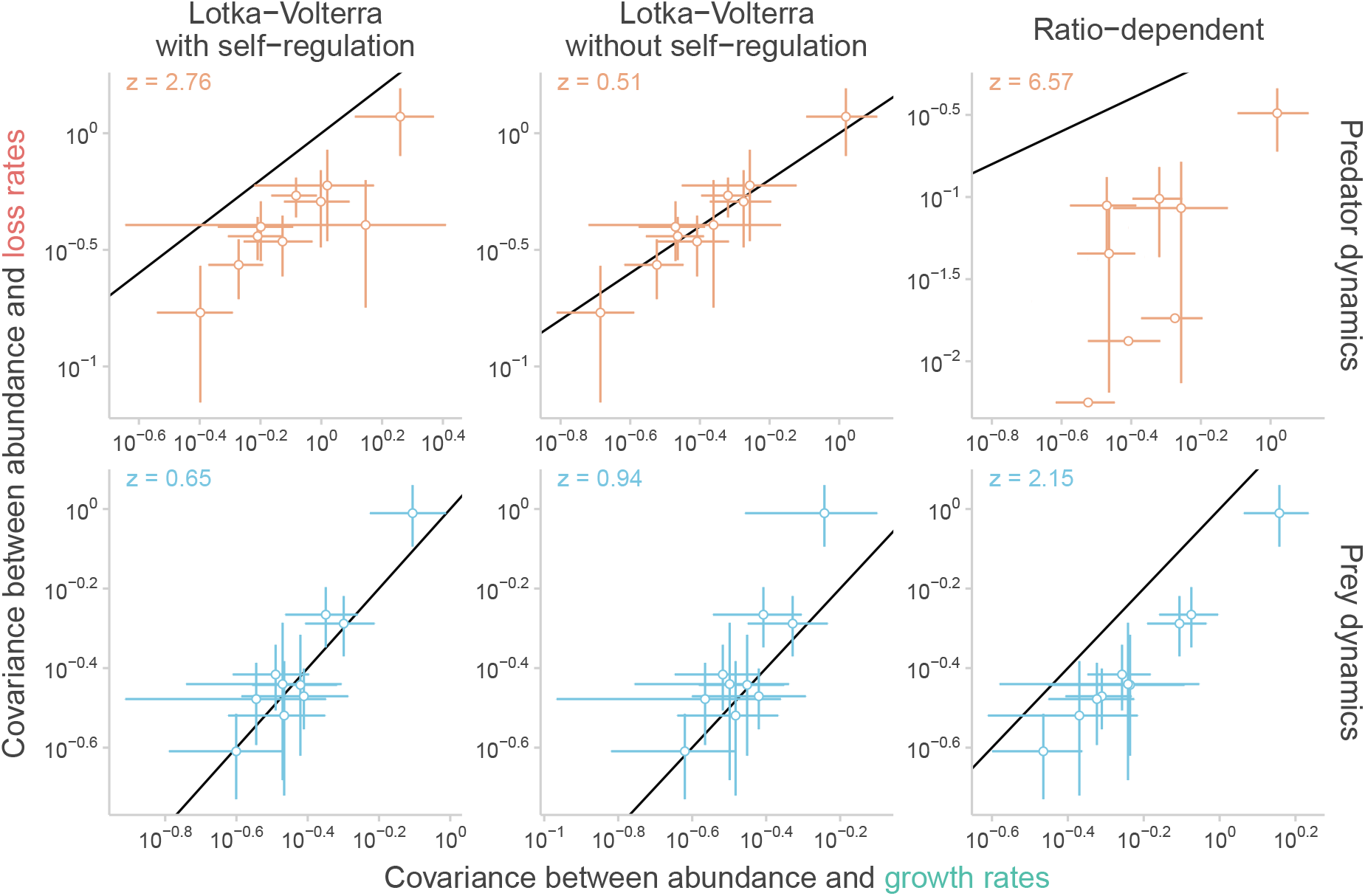
Reverse engineering the nature of predation. We apply the covariance criteria to study the functional form of the predation dynamics. We analyze the dataset from Blasius, Rudolf, et al. (2020) with 10 replicates of an experiment studying the aquatic invertebrate *Brachionus calyciflorus* (prey) and the green algae *Monoraphidium minutum* (predator) under varying conditions. Rows represent either prey (blue) or predator (orange) dynamics, while columns compare three models: Lotka-Volterra with self-regulation, Lotka-Volterra without self-regulation, and a ratio-dependent model. Each panel compares normalized covariances between gain/loss rates and abundance (x and y-axis, respectively), and the diagonal line denotes where the two covariances are equal. Each data point represents a replicate, and the error bars depict the 95% confidence interval. The value in the upper-left corner of each panel displays the average z-score of the replicates within that panel. We find that the Lotka-Volterra model with self-regulation best captures prey dynamics, while the Lotka-Volterra model without self-regulation best describes predator dynamics (see Figure S2 for further statistical details). These findings suggest that a prey-dependent functional form, as used in the Lotka-Volterra model, is more appropriate to describe predation in this system compared to a ratio-dependent model.

In addition, our analysis sheds light on a long-standing question about where self-regulation emerges in predator prey systems. We find that self-regulation may play a role in the dynamics of the prey species, but is not supported in the predator species. This observation is consistent with the broader ecological hypothesis that top predators lack strong self-regulating mechanisms (Pimm and Lawton, 1977; Tilman, 1982; Chesson, 2012; Song and Saavedra, 2021). One consequence of self-regulation in the prey species, as found here, is an implied role for stochasticity in shaping the persistent cycles characteristic of predator-prey systems. Without stochasticity, self-regulation within the prey population drives the system towards a stable equilibrium (Murdoch, Briggs, and Nisbet, 2013). However, when this self regulation interacts with environmental stochasticity, the equilibrium is disrupted and transient dynamics can cause indefinite fluctuations(Gurney and Nisbet, 1978; Nisbet and Gurney, 2003).

### 3.2 Dissecting ecological and evolutionary processes

Evolutionary and ecological processes can operate on similar timescales (Schoener, 2011; Loreau, Jarne, and Martiny, 2023). Prey-predator dynamics, in particular, have emerged as a prime example of such rapid evolution (Yoshida, Jones, et al., 2003; Mittelbach and McGill, 2019). However, a major modeling challenge lies in determining where in the ecological system to incorporate evolution: should we focus on prey evolution (Marrow, Law, and Cannings, 1992; Abrams, 1997), predator evolution (Abrams, 1992), or their simultaneous co-evolution (Dieckmann and Law, 1996; Gavrilets, 1997). Unfortunately, multiple models, each incorporating different assumptions about which species evolve, can produce similar observable patterns, including for example the synchrony of predator and prey population cycles (Abrams, 2000). This makes it difficult to pinpoint the specific evolutionary processes operating within the interaction based solely on qualitative observations of the data. Fortunately, the covariance criteria, with its ability to test how well a model captures the key dynamics of *each* species, offers a promising avenue to pinpoint the specific evolutionary processes at play.

To examine how evolution shapes the dynamics of predator and prey, we must first establish a baseline: how do prey-predator dynamics appear without rapid evolution? Building on our earlier finding (Figure 3), we propose the LV model with self-regulation for the prey and without self-regulation for the predator as a candidate. To test this model’s validity, we analyzed 18 time series across diverse ecosystems where rapid evolution is not thought to be operating in a major way (Utida, 1957; Huffaker et al., 1958; Barnet, Daft, and Stewart, 1981; Dulos and Marchand, 1984; Luckinbill, 1973; Luckinbill, 1974; Veilleux, 1976; Blasius, Rudolf, et al., 2020), compiled and processed by Hiltunen et al. (2014) except for Blasius, Rudolf, et al. (2020) (Figure S3). Applying the covariance criteria to these datasets, we find that the proposed form of the LV model generally describes both prey and predator dynamics well in the absence of rapid evolution (Figure 4A-B, statistical analysis in Figure S4).

**Figure 4.**
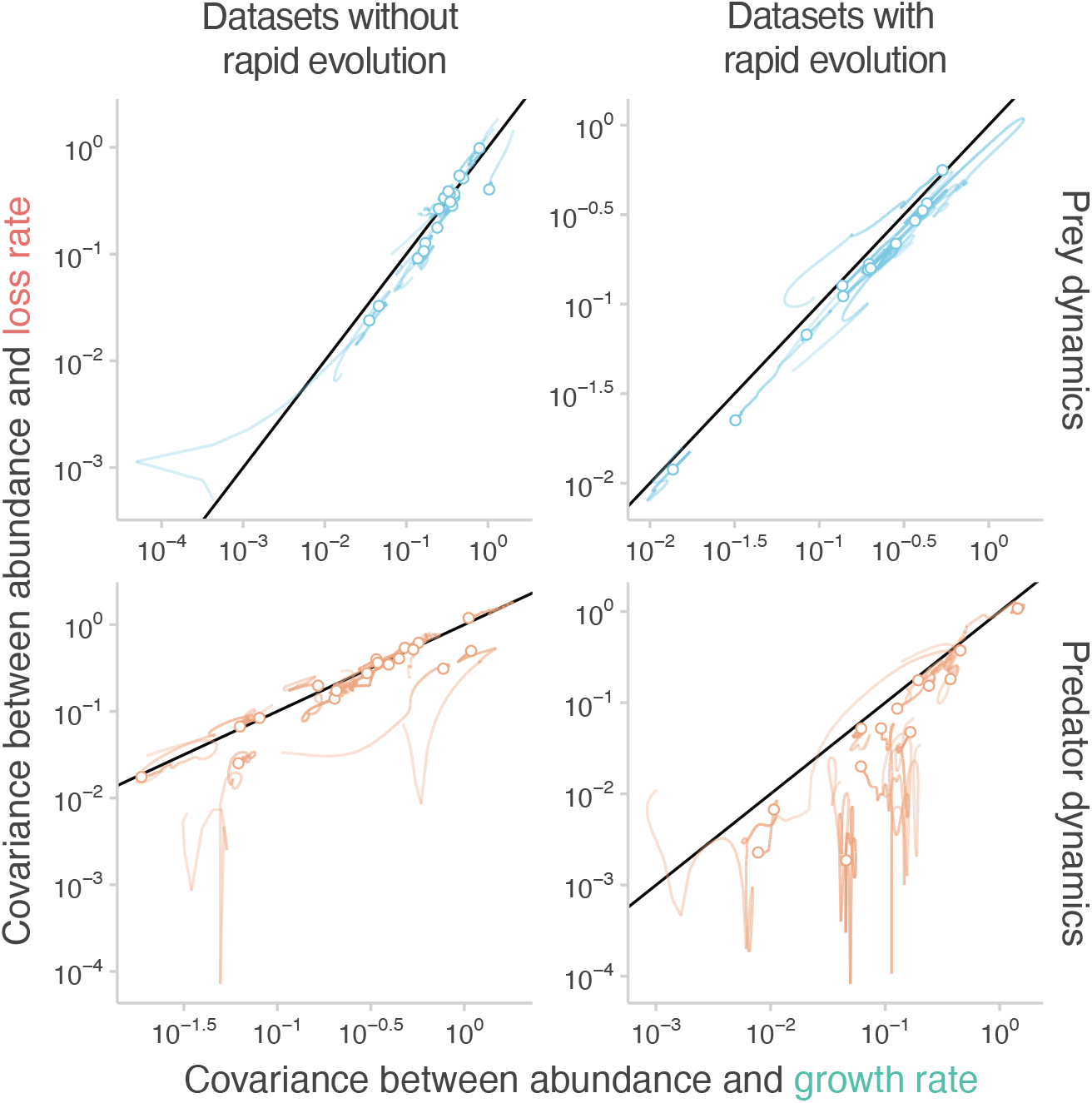
Dissecting ecological and evolutionary processes. We apply the covariance criteria to study how rapid evolution affects population dynamics in predator-prey systems. Inspired by previous analyses, we evaluate the Lotka-Volterra (LV) model with self-regulating prey. Rows represent either prey (blue) or predator (orange) dynamics. Columns differentiate between datasets with (right panels; 13 datasets) or without (left panels; 18 datasets) evidence of rapid evolution. Each panel compares normalized covariances between gain/loss rates and abundance (x and y-axis, respectively), and the diagonal line denotes where the two covariances are equal. Each line represents the results of the covariance criteria test applied to a specific dataset, using a different time window within that dataset. The transparency of the line indicates the size of the time window used: less transparent lines signify larger time windows, while a dot represents the analysis using the full time range of the dataset. We find that, without rapid evolution (left panels), the LV model effectively describes both prey and predator dynamics across ecosystems. In contrast, with rapid evolution (right panels), the LV model remains suitable for prey dynamics but not for predator dynamics. See Figure S4 for further statistical details. These results guide how to incorporate rapid evolution in modeling prey-predator dynamics.

With a reliable “no evolution” baseline model in hand, we can now ask: how does rapid evolution reshape the covariance structure of the system—in the prey, the predator, or both? To address this question, we analyzed 13 prey-predator time series where rapid evolution is empirically observed (Utida, 1957; Tsuchiya et al., 1972; Jost et al., 1973; Canale et al., 1973; Van den Ende, 1973; Veilleux, 1976; Boraas, 1980; Bohannan and Lenski, 1997; Yoshida, Jones, et al., 2003; Yoshida, Ellner, et al., 2007). These datasets, compiled and processed by Hiltunen et al. (2014), encompass a diverse range of ecosystems, providing an ideal testbed. The covariance criteria reveal a striking pattern: Predator species exhibit significant deviations from the baseline LV model (with no rapid evolution), suggesting the LV model no longer holds (Figure 4D and Figure S4). In contrast, prey species continue to adhere to the LV model’s predictions (Figure 4C and Figure S4).

These findings suggest that incorporating rapid evolution might require modifications to the predator component of the LV model, but likely not the prey component. A caveat, though, is that we cannot pinpoint specific evolutionary mechanisms as we have exclusively focused on phenomenological models. It is possible that evolution occurs in the predator’s capture-related traits and/or the prey’s defensive traits, but phenomenologically, only the predator seems to respond to these evolutionary changes in one or both species. Another caveat is that prey species may have different intrinsic gain rates with or without rapid evolution. Due to the non-parametric nature of the covariance criteria test, we cannot detect these potential differences because they share the same model structure. Despite these limitations, our findings provide guidance for selecting current eco-evolutionary models and catalyzing the development of new ones.

### 3.3 Detecting signals of higher-order interactions

Higher-order interactions (HOIs), where a third species modifies interactions between a pair, have long fascinated ecologists (Vandermeer, 1969; Case and Bender, 1981). Yet, detecting their existence remains challenging. Experimental manipulations, while ideal, are often logistically difficult (Mickalide and Kuehn, 2019; Barbosa, Fernandes, and Morris, 2023). A common alternative is to infer HOIs through model fitting (Mayfield and Stouffer, 2017; Li et al., 2021; Lai et al., 2022). However, since HOIs introduce more parameters, models can overfit the data, giving the illusion of HOIs where none exist (Dyson et al., 2004; Mayer, Khairy, and Howard, 2010). While regularization methods and information criteria can mitigate this issue (Tredennick et al., 2021; Aho, Derryberry, and Peterson, 2014), biases may still persist.

In contrast, the covariance criteria offer a compelling alternative for detecting potential HOIs, as they are inherently less susceptible to overfitting. Specifically, HOIs, when encoded in a model, change the predicted covariance structure. If that model was applied to a dataset with no true HOIs, a mismatch between the model’s predictions and the observed data would emerge. To demonstrate this, we analyze a high-quality, long-term dataset of a rocky intertidal community in Goat Island Bay, New Zealand (Benincà et al., 2015). This dataset tracks the monthly percent cover of barnacles, mussels, and algae for over 20 years. Benincà et al. (2015) proposed a model without HOIs for mussel dynamics:

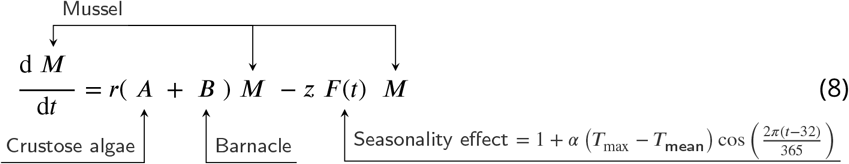

where ***M*** is the cover of mussels, ***B*** is the cover of barnacles, ***A*** is the cover of crustose algae, *r* is the rate at which area covered by those two species is colonized by mussels, *z* is the constant death rate of mussels, and *F* (*t*) represents the effects of seasonality, which is a complex function of abiotic factors.

Despite the complexity of this model, it has a simple covariance structure

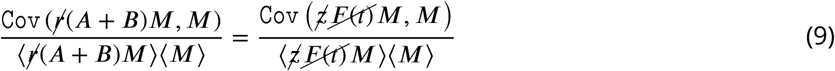

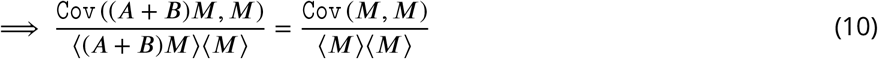

We can cancel the mussel colonization (*r*) and death rate (*z*) is because they are constant, and can cancel the effects of seasonality ***F*** (*t*) because ***F*** (*t*) is independent of the fluctuations of mussels ***M*** (p value = 0.72 with nonlinear correlation test in Chatterjee (2021)).

Additionally, we considered two further models. One model assumes mussel growth depends only on a HOI—the interactive effect of algae and barnacles on mussel colonization:

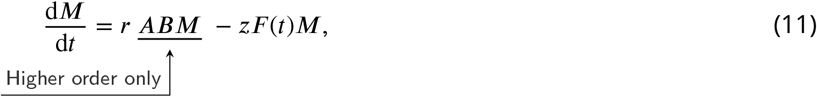

and the other model combines the pairwise and higher order interactions:

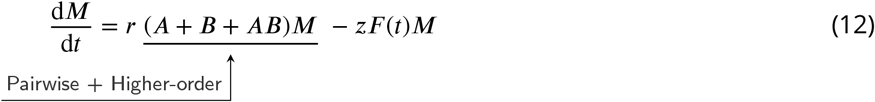

We then test the covariance structure from all three models above (Eqns. 8, 11, and 12) against the empirical data (Figure 5). We find that the HOI-only model (Eqn. 11) fails the test entirely (z-score = 6.17, gain rate covariance with density is quite different from the loss rate covariance). In contrast, the pairwise only model (Eqn. 8) shows similar covariances for the gain and loss rates (z-score = 1.75). The pairwise + higher-order interaction model (Eqn. 12) almost perfectly explains the data (z-score = 0.09), suggesting the presence of HOIs within this system.

**Figure 5.**
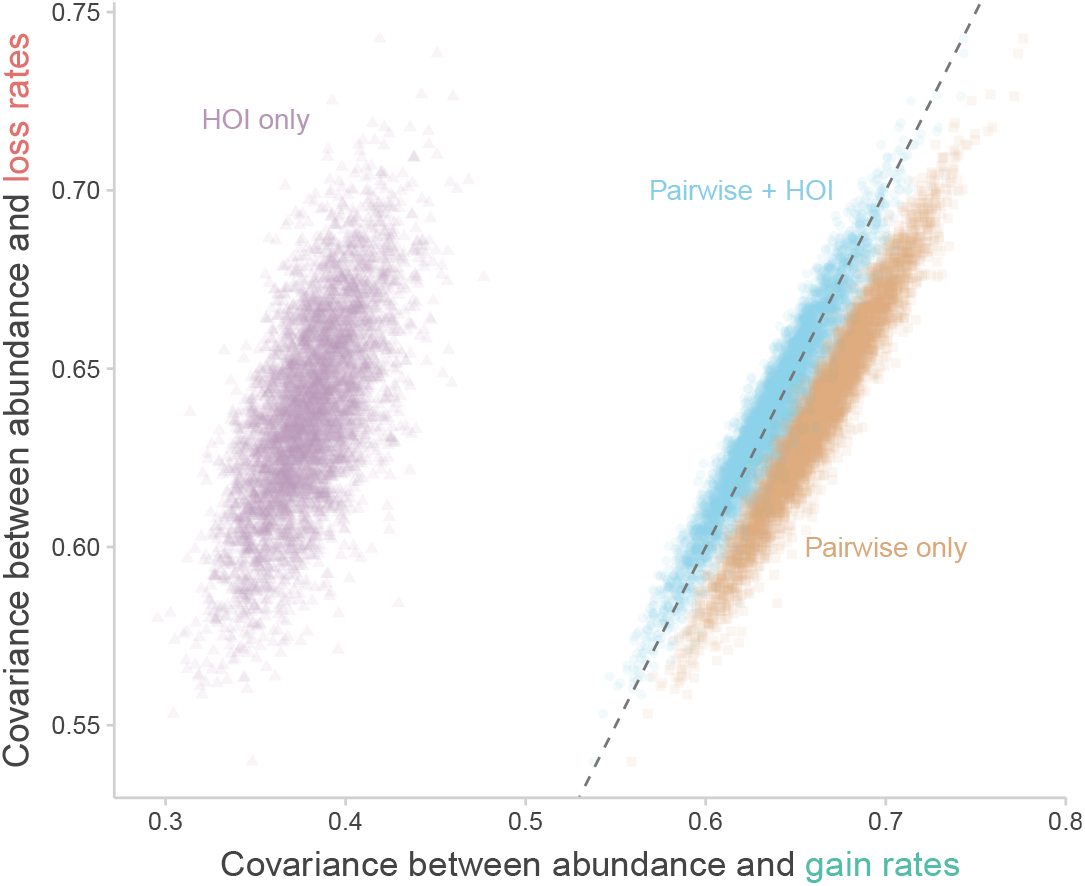
Detecting signals of higher-order interactions. We apply the covariance criteria to study the presence of higher-order interactions in a rocky intertidal community (Benincà et al., 2015). Specifically, whether crustose algae and barnacles interact with mussels exclusively through pairwise interactions or whether a higher-order interaction is present. Three models are evaluated: pairwise interactions only (orange), higher-order interaction only (purple), and a combination of both (blue). The x-axis represents the covariance between abundance and gain rates, while the y-axis represents the covariance between abundance and loss rates. Points are derived from 1000 bootstrapping replicates. The higher-order only model (purple) shows a significant mismatch in covariance values, indicating its inadequacy. The pairwise interaction model (orange) aligns more closely but still deviates statistically from the observed loss covariance. In contrast, the model incorporating both pairwise and higher-order interactions (blue) accurately captures the loss covariance. Figure S6 shows further statistical analysis. These findings strongly suggest that both pairwise and higher-order interactions between crustose algae, barnacles, and mussels play a significant role in influencing mussel dynamics within this community.

## 4 Discussion

We introduce the covariance criteria as a powerful, assumption-light framework for validating ecological models against time series data. The key insight is that every dynamical model imposes unique constraints on the permissible covariance structures relating population abundances, gain rates, and loss rates. If the empirical data satisfy these constraints, we can be confident the model provides a useful approximation capturing core aspects of the system’s dynamics. Conversely, violations of the covariance criteria provide quantitative evidence that the model is fundamentally inadequate, at least for the particular species and conditions examined.

Theoretically, the covariance criteria exhibit remarkable generality, applying across ecological dynamics ranging from simple equilibria to complex non-equilibrium systems with non-Markovian delays and external stochastic forcing. Computationally, the criteria are efficient to evaluate and often operate non-parametrically, eliminating the need to specify all model parameters from data. Perhaps most crucially from an empirical standpoint, the covariance criteria can be readily applied to the limited and noisy time series data common in ecological studies. As demonstrated through our three case studies, this approach consistently supports ecological models aligning with prevailing ecological understanding, while decisively rejecting those failing to capture underlying dynamics. In an era of rapidly accumulating high-quality ecological data, this approach subjects theorists’ ideas to rigorous scrutiny, and facilitates a better dialogue between ecological theory and empirical reality.

Theoretical ecologists have often been criticized for validating models with a low bar for consistency with data (Rykiel Jr, 1996; Bascompte, Jordano, and Olesen, 2006; Holland, Okuyama, and DeAngelis, 2006). The low validation bar allows a multiplicity of models to appear acceptable, even when their predicted mechanisms are vastly different, leading to insufficient confidence in any particular model. However, this raises a question: if we set a more rigorous quantitative bar, would all ecological models fail? This concern may explain the limited attention the covariance criteria has received beyond its originators (Joly-Smith, Wang, and Hilfinger, 2021; Wittenstein, Leibovich, and Hilfinger, 2022; Joly-Smith, Talpur, et al., 2023). After all, as Robert May wrote, “the models of biological communities tend rather to be of a very general, strategic kind” (May, 2002). We initially expected most ecological models to struggle to meet the strict covariance criteria. Much to our surprise, however, we found that the classic Lotka-Volterra model withstood the test across a wide range of consumer-resource systems. This stands in direct contrast to the common perception, echoed in many introductory ecology texts (Morin, 2009; Mittelbach and McGill, 2019), that the LV model is overly simplistic and misses crucial biological details. Our findings could help explain the recent success of the LV model in predicting some ecological patterns (Barbier et al., 2021; Hu et al., 2022; Père, Terenzi, and Werner, 2024).

The covariance criteria represents a fundamentally different approach to model validation than machine learning methods like symbolic regression (Chen, Angulo, and Liu, 2019; Martin, Munch, and Hein, 2018; Cardoso et al., 2020). Those machine learning techniques aim to directly distill mathematical models from patterns in empirical data, with minimal a priori knowledge of the system. In contrast, the covariance criteria retains theory-based model building at its core, as it begins with a theorist-proposed dynamical model inspired by natural history. Importantly, covariance criteria and machine learning approaches can work together. For example, they could be combined into a synergistic modeling pipeline where machine learning suggests new model structures, theorists use their expertise to explain the mechanisms, and the covariance criteria rigorously tests the resulting models against new data.

The covariance criteria could be useful for evaluating more than just population dynamics over time, as we focussed on here. For example, the criteria could be used to test models for how population abundances vary over space, analogous to common ecological approaches of substituting space for time when aiming to understand long term dynamics (Pickett, 1989; Wogan and Wang, 2018). However, doing so would require making additional assumptions about how model parameters vary across locations. Future work could also adapt the criteria to model the dynamics of other types of empirical data, such as temporal changes in trait values or single nucleotide polymorphisms.

Despite its advantages, the covariance criteria have limitations. It is most effective when a species has only few direct interactions with other species. This is because the criteria partitions the model into gain and loss components without dissecting their underlying process in any detail. When species interact directly with many rather than a few species, the gain and loss terms can become highly complex, potentially requiring the estimation of additional parameters. This increased parameterization makes the analysis more complex and diminishes the advantage of the covariance criteria being non-parametric in simpler cases. In practice, however, this may not be a major limitation because, outside of microbiome data analyzed with high-throughput sequencing, few long-term time series include a large number of interacting species.

As ecology grapples with increasingly complex challenges, from climate change to biodiversity loss, the need for reliable models has never been greater. The covariance criteria approach offers a path toward greater confidence in our ecological understanding by rigorously testing models across a wide range of problems. The approach could help validate models of species range shifts under climate change, or models of food web robustness to species invasions and extinctions. By providing a more rigorous foundation for model validation, we hope this method can contribute to more accurate predictions of ecosystem responses to environmental perturbations and more effective conservation strategies.

## Data availability

Empirical data of aquatic invertebrate and the green algae is available from doi.org/10.1038/s41586-019-1857-0. Empirical dataset of consumer-resource dynamics is available from doi.org/10.1111/ele.12291. The R package ecoModelOracle to run the analysis is available on GitHub at github.com/clsong/ecoModelOracle.

